# *Porphyromonas gingivalis* outer membrane vesicles alter neuronal architecture and Tau phosphorylation in the embryonic mouse brain

**DOI:** 10.1101/2024.09.03.611094

**Authors:** Adrienne Bradley, Lauren Mashburn-Warren, Lexie C. Blalock, Francesca Scarpetti, Christian L. Lauber

## Abstract

*Porphyromonas gingivalis* (Pg) is an oral bacterial pathogen that has been associated with systemic inflammation and adverse pregnancy outcomes such as low birth weight and preterm birth. Pg drives these sequelae through virulence factors decorating the outer membrane that are present on non-replicative outer membrane vesicles (OMV) that are suspected to be transmitted systemically. Given that Pg abundance can increase during pregnancy, it is not well known whether Pg-OMV can have deleterious effects on the brain of the developing fetus. We tested this possibility by treating pregnant C57/Bl6 mice with PBS (control) and OMV from ATCC 33277 by tail vein injection every other day from gestational age 3 to 17. At gestational age 18.5, we measured dam and pup weights and collected pup brains to quantify changes in inflammation, cortical neuron density, and Tau phosphorylated at Thr231. Dam and pup weights were not altered by Pg-OMV exposure, but pup brain weight was significantly decreased in the Pg-OMV treatment group. We found a significant increase of Iba-1, indicative of microglia activation, although the overall levels of IL-1β, IL-6, TNFα, IL-4, IL-10, and TGFβ mRNA transcripts were not different between the treatment groups. Differences in IL-1β, IL-6, and TNFα concentrations by ELISA showed IL-6 was significantly lower in Pg-OMV brains. Cortical neuron density was modified by treatment with Pg-OMV as immunofluorescence showed significant decreases in Cux1 and SatB2. Overall Thr231 was increased in pups exposed to Pg-OMV with the appearance of a secondary band of 60 kD. Together these results demonstrate that Pg-OMV can significantly modify the embryonic brain and suggests that Pg may impact offspring development via multiple mechanisms.

## Introduction

Pregnancy induces significant changes in mothers including shifts in the abundance of oral bacteria (1-4). Increased blood flow and pregnancy related hormones allows for the growth of pathogens like *Porphyromonas gingivalis* (Pg) an organism associated with oral infections that are increasingly linked to adverse pregnancy and neurodevelopmental outcomes (5-7). Increased Pg presence is estimated to be prevalent in 40% of pregnancies in the US and 2-11% worldwide (8, 9). This presence of bacteria sometimes resulting in periodontitis, a significant dental infection, can drive systemic inflammation (10-12). Preterm birth, preeclampsia, and low birth weight are well-characterized effects of maternal Pg infections and Pg DNA has been detected in the placenta and amnionic fluid. Pg driven outcomes affecting children have focused on pregnancy outcomes (13-18) and only recently has Pg received attention as a modulator of fetal development and remains an understudied host-pathogen interaction (19, 20).

*P. gingivalis* has been extensively studied to determine its role in modifying pregnancy. Rodent models of Pg infection demonstrate that exposure to Pg can result in systemic inflammation, low birth weight and pre-term delivery (21-25). Pg mediates these and other effects thorough a number of virulence factors that modify host responses including lipopolysaccharide (LPS) and proteases known as gingipains (26-32). Both are also present on the surface of OMV that are now suspected to be transmitted systemically in the blood stream (20, 32, 33). LPS and the gingipains interact with the immune system but are unique compared to other gram-negative bacteria. For instance, Pg LPS is less immunogenic than *Escherichia coli* LPS but also causes inflammation and the production of pro-inflammatory cytokines (34, 35). The gingipains are unique to Pg and they modify cytokine expression that allows Pg to evade the host immune response and establish periodontal infections that drive systemic inflammation (36-39). Pg infection also modifies the placenta to cause spiral artery remodeling that can result in a hypoxic state linked to preeclampsia and fetal growth restriction (40). However, whether Pg-OMV may induce fetal growth restriction—a common outcome in animal models and humans with Pg infections of clinical significance—is not yet known.

*P. gingivalis* is associated with maternal inflammation and adverse pregnancy outcomes that increase the risk of modified developmental trajectories of offspring (17, 41). Live Pg is well demonstrated to increase inflammation in animal models and exposure to Pg-OMV recapitulates these observations as the vesicles harbor the same virulence factors. Ishida et al. reported increased Iba-1 indicative of microglia activation and modified neuroarchitecture in E 20 pup brains from dams with an established Pg infection prior to pregnancy (19). These alterations were accompanied by declines in cognitive ability, and it was suggested the combined effects on the embryonic brain resulted in neuroinflammation that drove these changes. Similarly, Gong et al. described a neuroinflammatory phenotype in adult mice where exposure to Pg-OMV for 3 months increased IL-1β and NLRP3 expression concurrent with modification of the neuroarchitecture and behavior (33). Together these studies suggest that long-term exposure to Pg and Pg-OMV can result in immune phenotypes indicative of a pro-inflammatory response that would be expected from activated microglia in the M1 activation state (42, 43). However, the maternal immune system functions differently than the non-pregnant state and changes in inflammation and neuroarchitecture related to pathogen exposure *in utero* after pregnancy is established may also differ. This knowledge gap limits our efforts to refine predictions of the potential impact of pathogen-driven inflammation on the developmental trajectory of offspring.

Tau is an essential protein with multiple roles in maintaining neuron homeostasis (44-46). Proper function is dependent on phosphorylation that can occur at multiple sites and can be modified by inflammation and exposure to pathogens such as Pg (47-49). Phosphorylated Tau (p-Tau) has less affinity for tubulins and self-aggregates into β-amyloid plaques that are a hallmark of neuroinflammatory diseases such as Alzheimer’s and Parkinson’s diseases in older adults (50). The resulting Tau tangles can trigger an immune response that is sustained through microglial NFκβ signaling and thus are considered pathogenic forms of the protein (51, 52). In a mouse model, Gong et al. showed that Pg-OMV alter the Tau profile and could result in β-amyloid plaques accompanied by increased inflammation (33). However, relatively less is known about the role of p-Tau in the developing brain and whether exposure to pathogens during pregnancy alters this profile. A survey of fetal and adult Tau phosphorylation in humans demonstrated significant overlap in p-Tau where some of the pathogenic forms of Tau in adults with Alzheimer’s disease were also prevalent in the fetal brain suggesting p-Tau in children does not have a pathogenic effect as observed in adults (53). Interestingly, Tau is hyperphosphorylated in the embryonic rodent brain but knock-down of Tau during gestation in mice can reduce neuronal migration to all cortical layers indicating that p-Tau is an important part of normal brain development in mice (47, 49, 54). Whether or not Pg-OMV may modulate Tau phosphorylation in developing brains *in utero* is an open question. Addressing this knowledge gap may provide valuable insight into the mechanisms by which Pg is able to modify developmental trajectories of children.

Herein, we describe a study to determine the effect of Pg-OMV on the developing mouse brain to understand this host-pathogen interaction that frequently occurs in the human population. To simulate a maternal infection with Pg, pregnant C57/Bl6 mice were exposed to a consistent dose of Pg-OMV via tail vein injection. At gestational age (GA) 18.5, inflammatory responses, concentration of Tau Thr231 (Thr231) and neuron density in the cortices of embryonic pup brains were assessed to augment the understanding of Pg-modulated brain development that may be relevant to pathogen exposure during pregnancy.

## Results

### Pup Body and Brain Weights

Previous studies in mice have demonstrated that dams with Pg infection or those consistently dosed with Pg bacteria during pregnancy can result in low birth weight of pups. We recorded weight of the dams, pups, and pup brains at GA 18.5 (Fig 1A) to determine whether exposure to Pg-OMV would have the same effect. Dam and pup weights were not significantly different in PBS and Pg-OMV groups (Fig 1B and 1C) but there was a significant decrease in the pup brain weight after 14 days of Pg-OMV treatment (Fig 1D, Mann-Whitney U p = 0.023,). We included an assessment of Hif1a as Pg is known to cause hypoxia in mouse models that is related to fetal growth restriction. Relative expression of HiF1a in the embryonic mouse brain was not significantly different between the control and treated groups indicating the OMV did not induce a hypoxic state *in utero* that could account for the decreased pup brain weights (Fig 1E).

**Fig 1.**
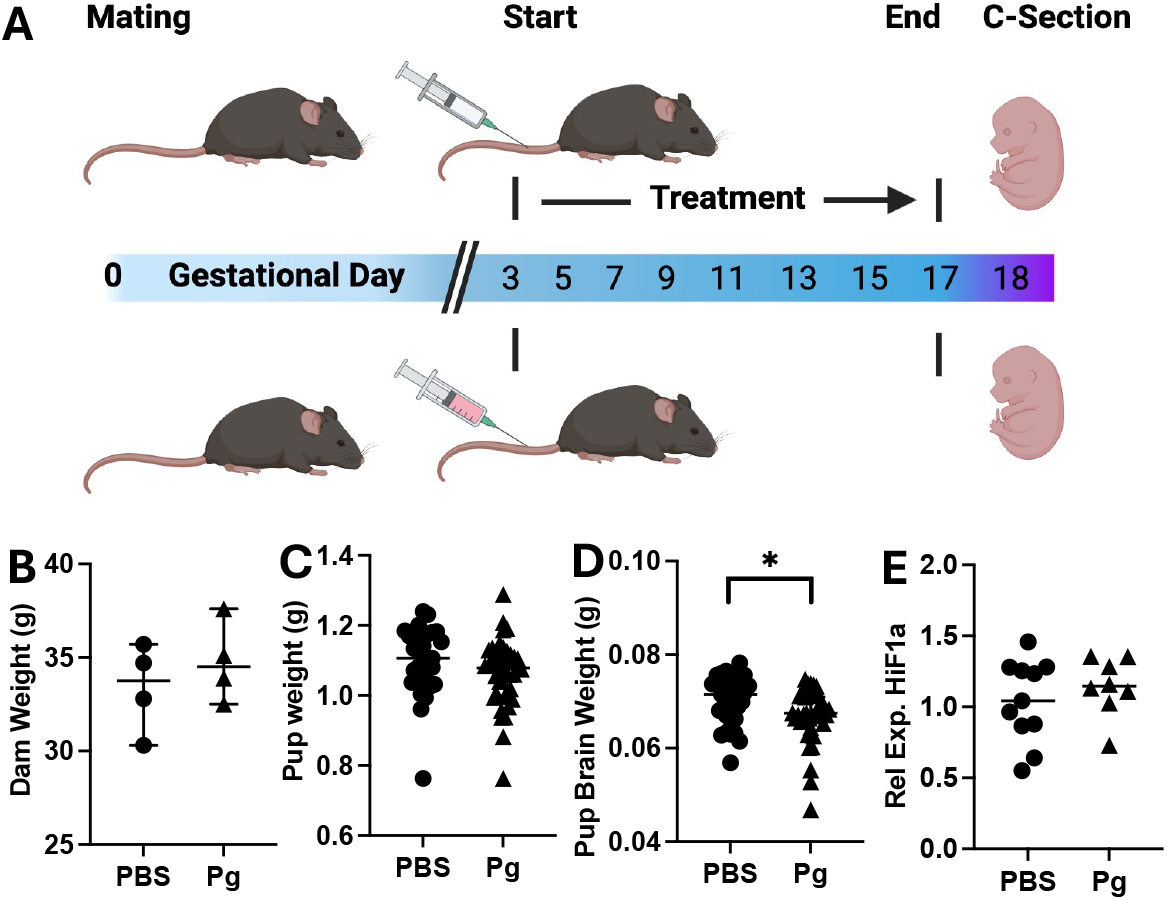
Experimental design and pup body and brain weights. (A) C57/Bl6 mice were mated for up to 3 days until the appearance of vaginal plugs. The first observation of a plug was considered GA 0. Female mice were separated from the males and treated with PBS or 50 ug Pg-OMV every other day from GA 3 to GA17. Dams were sacrificed on GA 18.5 and pups delivered by C-Section for tissue collection. (B) and (C) Weight of dams and pups were not significantly different from PBS controls. (D) Brain weight decreased significantly with Pg exposure (Mann-Whitney U p = 0.023). (E) Relative expression of Hif1a as indicator of hypoxia in the pup brain. Relative expression calculated by the 2^^-(ΔΔCt)^ method with GAPDH as the reference gene. Data are shown as median with 95% CI.

### Inflammation

Maternal inflammation can affect the developing embryo in mouse models, and we tested whether exposure to Pg-OMV during pregnancy would result in an inflammatory response concordant with increased presence of pathogens experienced during pregnancy. First, we assessed the concentration of Iba-1 as a surrogate of microglia activation and established this immune cell typically involved in pathogen defense was active in the embryonic mouse brain. Western blotting of whole brain homogenates showed that Iba-1 was significantly increased in Pg-OMV pup brains compared to PBS controls (Fig 2A and 2B, Mann-Whitney U p = 0.0003). Next, we tested whether increased Iba-1 was accompanied by increased concentration and expression of pro-inflammatory cytokines IL-1β, IL-6 and TNFα indicative of M1 activation that produces a pro-inflammatory response. ELISA quantification showed that PBS pups typically had higher levels of cytokines but only IL-6 was significantly greater in the PBS brains compared to pups from the Pg-OMV group (Fig 3D, Student’s T-test p = 0.002). The levels of IL-1β and TNFα showed similar trends but were not statistically different between the treatment groups (Fig 3A, 3D, 3G, p > 0.05 in all cases). We extended this analysis to include effects that may be apparent at the level of expression in the embryonic brain tissue. RT-qPCR analysis of these cytokines showed no significant difference between the PBS and Pg-OMV pups. Then, we tested the possibility that increased Iba-1 could result in an M2 or anti-inflammatory immune response given that microglia can be in two broad functional categories (42, 55). Analysis of IL-4, IL-10 and TGFβ typical of M2 activated microglia were not significantly different between the treatment groups (Fig 3 C, 3F, 3I, p > 0.05 in all cases). Lastly, given these results we tested whether the expression of NLRP3, MyD88 and NFκβ, which can be modified by exposure to Pg or it’s OMV, were also modified in our pups. RT-qPCR analysis of MyD88 and NLRP3 transcripts (Fig 4) had similar downward trends with Pg-OMV treatment but were not significantly different compared to controls (Fig 4A, and 4B, p > 0.05 in both cases). Interestingly, NFkβ transcripts that are typically reported as being upregulated by Pg were significantly decreased in the Pg-OMV group (Fig 4C, Student’s T-test p = 0.004).

**Fig 2.**
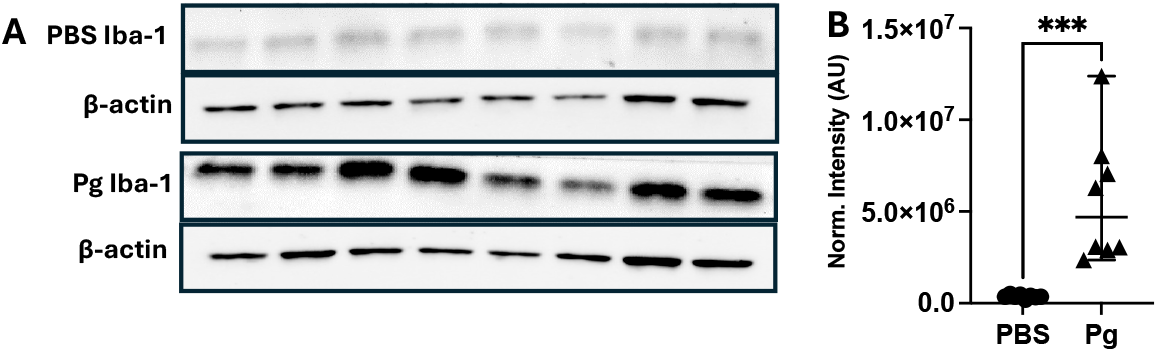
Increased Iba-1 in mouse pups exposed to Pg-OMV during pregnancy. **(A)** Western blot of Iba-1 and β-actin in GA 18.5 pups from PBS and Pg-OMV exposed dams. Full blot can be found in the Supporting information. **(B)** Comparison of band β-actin normalized signal intensity of Iba-1 showed significant increase in Iba-1 in the Pg-OMV group compared to controls (Mann-Whitney U p = 0.0003). Data are shown as median with 95% CI.

**Fig 3.**
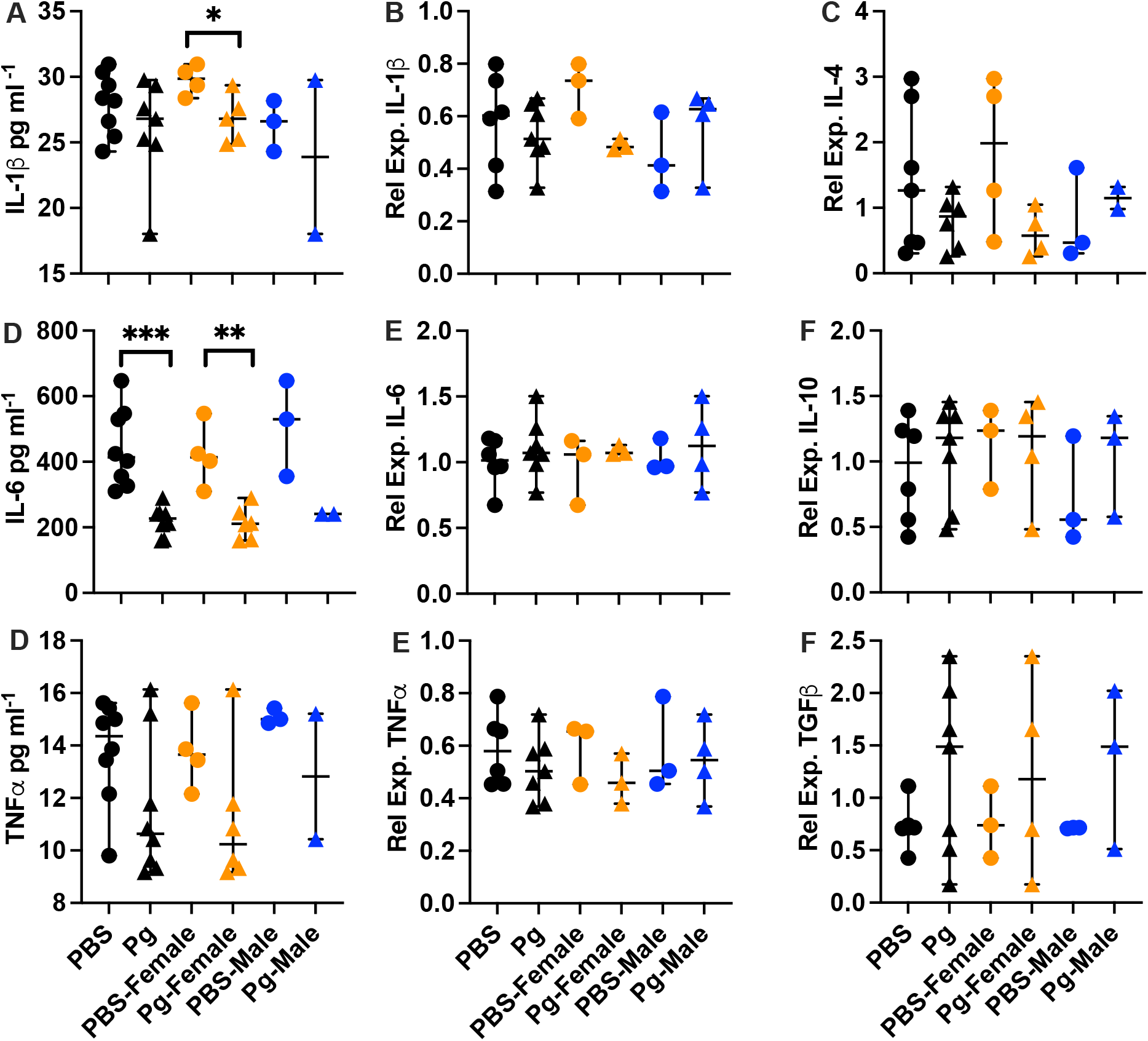
ELISA and RT-qPCR evaluation of cytokines in the embryonic mouse brain. **(A, D, G)**: ELISA of IL-1β, IL-6, and TNFα in pg ml^-1^. Cytokines were extracted from frozen whole brains and quantified. IL-1β was significantly greater in the PBS female pups compared to the female pups from the Pg-OMV group (Fig 3A, Mann-Whitney U test p = 0.04). IL-6 was significantly greater in the PBS group and in the female pups compared to the OMV treatment group (Fig 3D, Student’s T-test p = 0.002 and 0.0095, respectively.) **(B, E, H)**: RT-qPCR of proinflammatory cytokines IL-1β, IL-6, and TNFα. **(C, F, I)**: RT-qPCR of anti-inflammatory cytokines IL-4, IL-10, and TGFβ. Total RNA was extracted from frozen whole brains and reverse transcribed to cDNA. Relative expression calculated by the 2^^-(ΔΔCt)^ method with GAPDH as the reference gene. Group differences for the ELISAs on the males was not determined due to low sample size (n = 2). Only significant differences are shown. Data are presented with the median and 95% CI. Symbols: Circles = PBS, triangles = Pg; orange = females; blue = males.

**Fig 4.**
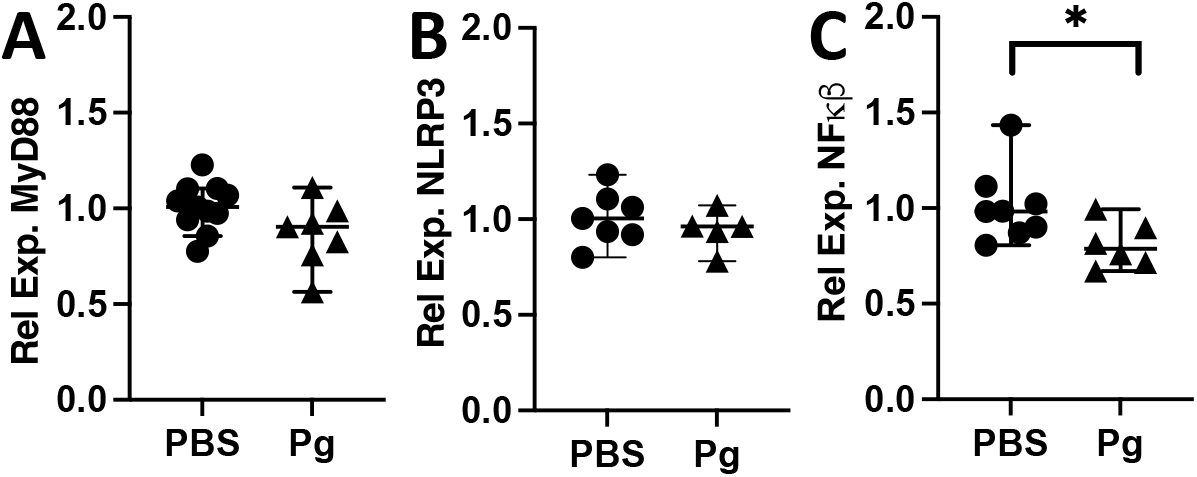
RT-qPCR evaluation of MYD88, NLRP3, and NFκβ in GA 18.5 pup brains. **(A)** MYD88 and **(B)** NLRP3 were not significantly different in pup brains from the control or treatment group. **(C)** NFkβ was significantly reduced in the pup brains (Student’s T-test p = 0.004). Total RNA was extracted from frozen whole brains and reverse transcribed to cDNA. Relative expression calculated by the 2^^-(ΔΔCt)^ method with GAPDH as the reference gene. Means across fore, mid, and hind brain sections are shown for each sample. Data are presented with the median value and 95% CI.

We were interested in identifying potential modifications in cytokine concentration and expression that may differ between female and male offspring as maternal inflammation may have sex specific effects. When parsed by sex, we observed that IL-6 and IL-1β concentrations were significantly lower in female pups from the Pg-OMV group compared to the female controls (Fig 3A, Mann-Whitney U test p = 0.04 and 3D, Student’s T-test p = 0.0095) and though TNFα was lower in Pg-female pups, it was not statistically different in our analysis (Fig 3A, 3D. 3G). The effect of Pg-OMV on transcription was highly variable for IL-1β, TNFα, IL-4, IL-10 and TGFβ but generally trended upwards (Fig 3B-3I) for the female pups. Trends in the males were similar but were not significant and likely result from low sample sizes (Fig 3), though we could not assess difference in cytokine concentrations as the Pg male group had only 2 observations (Fig 3A, 3D, 3G).

### Neuroarchitecture

Without a strong inflammatory phenotype, we tested the possibility that Pg-OMV treatment could nonetheless result in differences in the neuroarchitecture of the developing mouse brain. We examined the distribution of neuronal proteins Cux1, SatB2, and Ctip2 that are differentially expressed across the layers of the cortex in sagittal sections of embryonic mouse brains. The distribution and density of these proteins were modified by exposure to Pg-OMV compared to pups from the PBS group (Fig 5A and 5B). We saw significant decreases in Cux1 (Student’s T-test p = 0.03) and SatB2 (Students T-test p = 0.0001) signal intensity but not Ctip2 (Student’s T-test p = 0.064) although it also followed a similar downward trend in the Pg-OMV pup brains (Fig 5B). These findings suggests that cortical layering was drastically altered in Pg exposed animals.

**Fig 5.**
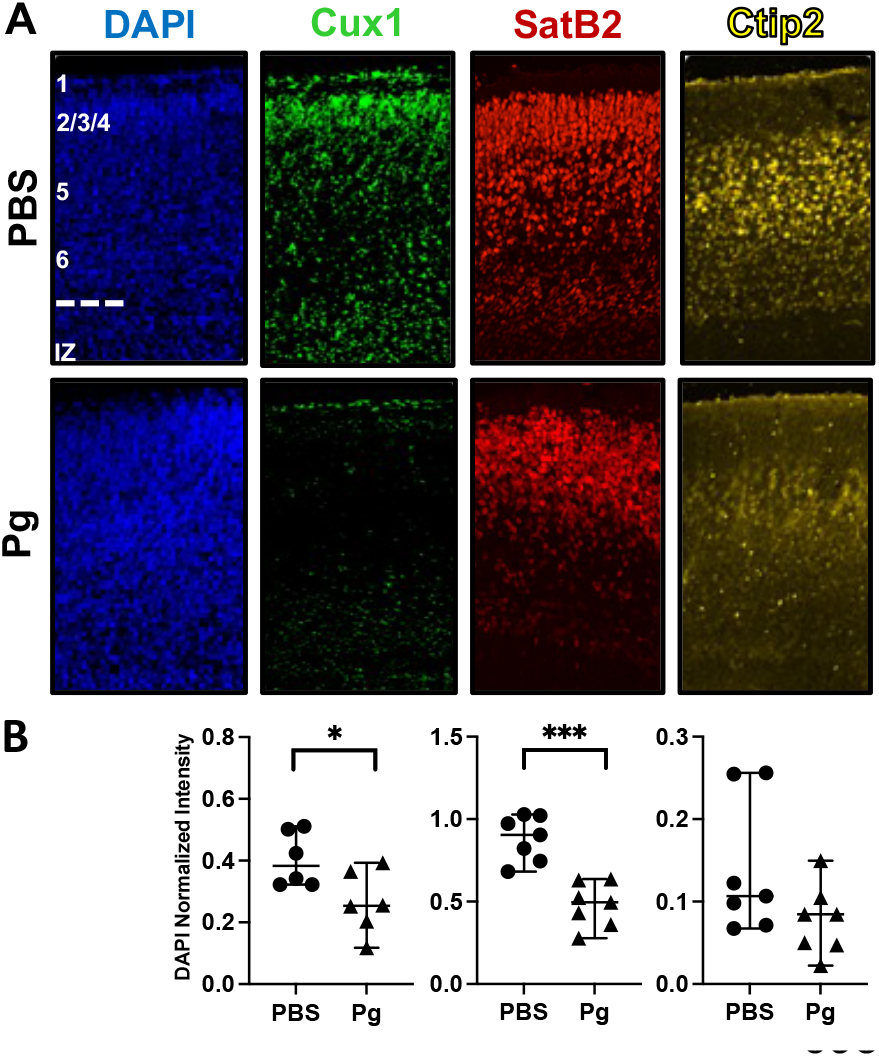
Modified intensity and location of cortical layer marker proteins following Pg-OMV exposure *in utero*. **(A)** Representative immunofluorescent images from sagittal sections of PFA fixed brains at GA 18.5 pup brains showing location of cortical layer markers Cux1, SatB2, and Ctip2. Approximate position of cortical layers 1-6 with the intraventricular zone (IZ) are shown in the PBS-DAPI image. **(B)** Quantification and comparison of cortical layer proteins stained with the marker antibodies. Intensities were averaged across three serial section per sample for statistical analysis. Cux1 (Student’s T-test p = 0.03) and SatB2 (students T-test p = 0.0001) were significantly reduced in the Pg-OMV group. Data are presented with the median and 95% CI.

### Tau Thr231

We evaluated whether Tau phosphorylated at Thr231 in GA 18.5 mouse brains was affected by exposure to Pg-OMV during gestation as this is one form of p-Tau implicated in neuroinflammatory diseases. Western blotting of proteins from whole brain homogenates showed Thr231 was present in PBS and Pg-OMV mouse brains at approximately 50 kD indicating that at this amnio acid, Tau was in a phosphorylated state concordant with other observations of Tau in the developing mouse brain (Fig 6A). We observed that an additional band at 60kD became more prominent in the Pg-OMV brains compared to controls (Fig 6A). Quantification of these bands showed that both the 50kD and 60kD bands were significantly greater in the brains from Pg-OMV exposed mice compared to the PBS control group (Fig 6B, 50 kD Student’s T-test p = 0.003, 60kD Student’s T-test p = 0.0006). The increased abundance of both (e.g. 50kD signal + 60kD signal = total Thr231) lead to significant overall increase of Thr231 in the Pg-OMV group (Fig 6B, total Thr231, Student’s T-test p = 0.001) indicating Pg-OMV may alter the total concentration and diversity of Thr231 isoforms during development in our mice.

**Fig 6.**
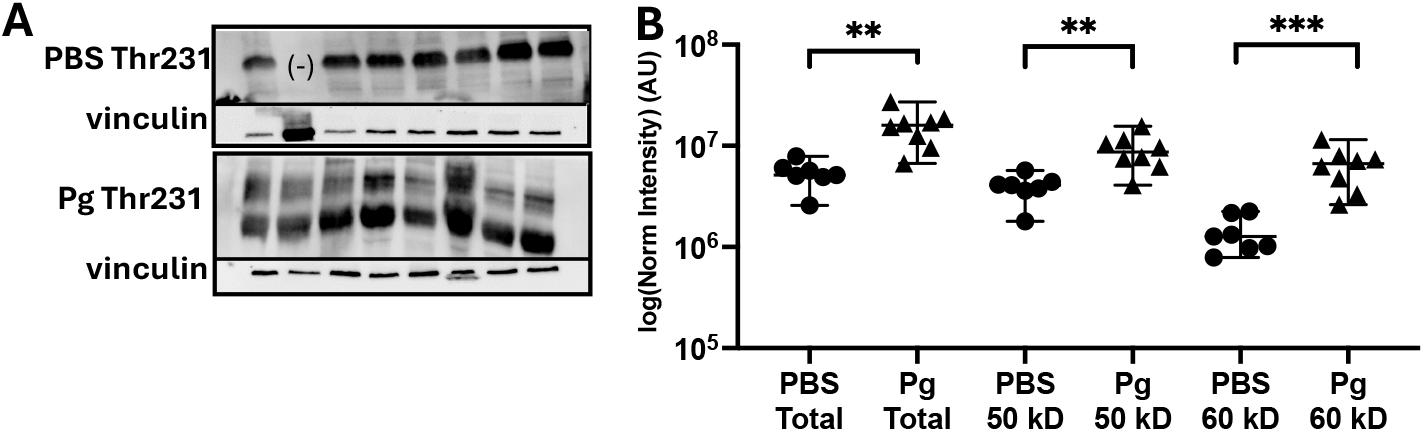
Thr231 is elevated in the embryonic mouse brain after exposure to Pg-OMV. **(A)** Images of Thr231 western blots. 20 ug of total protein extracted from whole, frozen GA 18.5 pup brains were analyzed. A band of approximately 60 kD became more prominent in the brains of all pups exposed to Pg-OMV during pregnancy. **(B)** Densitometry analysis of Thr231 abundance by western blot normalized to vinculin. Total Thr231 was estimated by summing the vinculin normalized intensity of the 50 kD + 60 kD bands. Total Thr231 (Student’s T-test p = 0.001), 50kD (Student’s T-test p = 0.003) and 60kD Thr231 quantities (Student’s T-test p = 0.0006) were significantly greater in the Pg-OMV brains. Data are presented with the median and 95% CI.

## Discussion

Animal models of Pg infection demonstrate modified pregnancy outcomes and can alter birth weight of offspring. Our experimental design did not recapitulate these observations as pup weights in the Pg-OMV group were not statistically different from PBS controls. Moreover, dam weights were also unchanged indicating that food and water intake were likely not different between the control and treatment groups. The mechanism behind reduced brain weight is not clear from our observations, although pup weight followed a similar downward trend suggesting that the effect of Pg-OMV was systemic for the developing embryos. It could be possible that exposure to pathogens after initiation of pregnancy is different from a state where the pathogen is already present and interacting with host defense mechanisms. Likewise, the dosage of Pg-OMV we used may have not been sufficient to recapitulate the full effect of Pg on pregnancy outcomes, where more robust responses of the host and fetus may be required to lower birth weights. This would coincide with unchanged expression of Hif1a. Nevertheless, treatment with a low dosage of Pg-OMV during pregnancy impacted the development of the embryonic brain that could have implications after birth.

*P. gingivalis* and Pg-OMV induced neuroinflammation is suspected to be involved in cognitive decline in aging cohorts and have a significant impact on development *in utero*. Our hypothesis that Pg-OMV would produce a pro-inflammatory response in the embryonic mouse brain was not observed. Cytokine concentrations and levels of mRNA transcripts in the pup brains tended to be lower in the Pg-OMV group. The lack of concordance with previous work was somewhat unexpected as most reports demonstrate systemic inflammation upon exposure to Pg or its OMV. Significant increases of Iba-1 in Pg-OMV exposed brains indicates sensing of bacterial-derived components, such as LPS, that can stimulate immune response, though this was not accompanied by a significant increase in cytokine production in our mice. Moreover, Pg has atypical LPS compared to organisms like *E. coli* LPS, that can bind to TLR-4, but can be recognized by alternative sensing pathways. We addressed this possibility by quantifying the expression of MYD88 and NLRP3 inflammasome mRNAs that are known to increase upon exposure to Pg or Pg-OMV. Our analysis found that concentrations of both were unaffected by OMV treatment in GA 18.5 pup brains and we posit several factors may be influencing our data. First, our results could originate from the delivery of Pg-OMV via tail vein injection which bypass sensing mechanisms such as dendritic cells of the gastrointestinal tract that would detect Pg and LPS being translocated from the oral cavity in saliva. Second, it is equally possible that Pg-OMV induced inflammation reported elsewhere may be an outcome in adult mice with long-term exposure to Pg or Pg-OMV but may not necessarily be an outcome for exposure after pregnancy has been established given the maternal immune system functions differently compared to the pre-pregnancy state. Thus, our observations may not be comparable to reports that use Pg infection or treatment with Pg or Pg-OMV at high doses over several weeks prior to inducing pregnancy. Importantly, the dosing scheme described here suggests there is a potential use of Pg-OMV as a proxy for bacteria to study effects on fetal tissues independent of systemic inflammation driven by common oral pathogens that become more prevalent during pregnancy. More work is needed to elucidate how the timing and dosage of pathogen exposure relative to the start of pregnancy modulates immune responses during gestation that could be associated with long-term consequences for offspring in human populations.

Several reports have quantified the impact of Pg and Pg-OMV on neurons as this cell type is affected by inflammation and is important in brain development. Our data indicate that Pg-OMV have a significant impact on neurons as their prevalence was modified in the cortical layers of the embryonic brain. Cux1, SatB2, and Ctip2 signals were lower overall and showed modified distribution in the Pg-OMV exposed brains compared to controls. The down regulation of SatB2 and Cux1 are particularly interesting as abnormal expression of both are associated with altered behavior in mice and are suspected to be a contributor to autism spectrum disorder-like symptoms in humans. Likewise, deficiencies in Cux1 expression have been linked to neurodevelopmental delays, while down regulation of Ctip2 is known to affect axonal growth and is involved in spiny neuron differentiation. The neuroarchitectural changes observed herein are aligned with previous studies, yet novel because these cortical layer aberrations were not accompanied by any indication of systemic inflammation. Collectively, these data suggest that Pg, or perhaps Pg-OMV specially, affect the developing brain through a different mechanism. Parsing out the effects of Pg-OMV on cell proliferation and migration appears to be critical in deciphering how Pg mediates effects on the developing brain and should be addressed in future studies to better understand this important host-pathogen interaction.

The importance of Tau in the developing brain has not been fully described but our observations indicate exposure to Pg-OMV can alter the amount of Thr231 in mouse embryos. Tauopathies have multiple effects on the central nervous system including sustained inflammation through NFκβ signaling microglia that detects the pathogenic form of Tau in adult animal models. Our data would appear to support the notion that Thr231 in the embryonic mouse brain is not pathogenic as an innate immune response, concordant with an inflammation-based hypothesis of neurodegenerative phenotypes, was not observed. Accordingly, we did not observe an increase in NFκβ that can encourage transcription of cytokines. We did not perform a comprehensive assessment of Tau phosphorylation sites, nor did we specifically address potential interactions between p-Tau and microglia. Nonetheless, the alteration of Thr231 abundance aids in explaining how changes in neuron migration and axon growth may result from exposure to Pg *in utero*; our observed changes in cortical layer markers could be the result of altered microtubule binding driven by altered p-Tau. Thus, much work remains to be done in elucidating the role of Pg and Pg-OMV on human neurodevelopment and cognitive trajectories from the early to elderly phases of life.

## Methods

### OMV Isolation

*Porphyromonas gingivalis* strain ATCC 33277 was grown in anaerobic conditions (85% N_2_, 10% CO_2_, 5% H_2_) using Tryptic Soy Broth (Difco cat# 211768) medium supplemented with 5.0 mg ml-1 hemin (Thermo Fisher, cat# AAA1116503), 1.0 mg ml^-1^ menadione (MP Biomedicals, cat # 102259), and 0.5 g L^-1^ L-cysteine-HC (MP Biomedicals, cat # 101446). A cleared supernatant was produced by pelleting cells at 8000 rpm for 15 minutes followed by filtration of the supernatant through a 0.45 um membrane filter under low vacuum pressure. Using Sorvall 11.5mL ultracentrifuge tubes and ultracrimp tube plugs, filtered supernatant was ultacentrifuged at 65,000 rpm for 1 hour at 4°C using a Sorvall WX Ultra 80 centrifuge. Immediately after centrifugation, supernatant was discarded and the pellet containing the OMV was resuspended in 500uL of PBS (56). The total protein of the OMV were quantified using the Pierce BCA assay with BSA as a standard and stored at -20°C for future use.

### Animal Breeding, Dosing and Tissue Collection

2-3 month old C57/Bl6 mice (male and female) were purchased from Jackson Labs (Maine, USA). All animals were housed in the animal research core under 12 h light/dark cycles. Mice were bred and dams were checked for plugs 2x/day. Once a plug was visible, the dam was removed and was marked as gestational day 0 (GA 0). Starting at E3, 50 µg Pg OMV were administered via tail vein injection every other day until GA 18.5. An equal volume of sterile PBS was used for the control group. Pregnant dams were anesthetized using isoflurane and pups were removed via C-sections. C Pups were decapitated and whole brain and placenta were collected from each pup. Half of the pup brains were randomly chosen and fixed using 4% paraformaldehyde followed by 30% sucrose prior to being mounted in O.C.T. and frozen for Immunohistochemical and Immunofluorescent analysis. The remaining brains were snap frozen in a liquid N_2_ at time of collection and used for protein, RNA, and DNA extraction. Fixed tissues were stored at 4°C and frozen tissues were stored at -80°C. All experimental procedures were approved by the IACUC under protocol AR21-00010.

### Protein Extraction, Western Blotting and ELISA

Protein from snap-frozen whole brains were extracted using RIPA Buffer (cell signaling) with protease inhibitor cocktail (ThermoFisher). Concentration was determined prior to loading using the Pierce BCA Assay Kit (ThermoFisher) with subsequent colorimetric detection using BioTek Synergy H1 microplate reader. Samples were denatured in Laemmli buffer at 95°C for 5 min. 50 µg total protein from each sample was loaded onto 10% Mini-PROTEAN® TGX™ Precast Protein Gels (Bio-Rad) and separated for 1 h at 150 V. Proteins were transferred onto nitrocellulose or PVDF membranes using Turbo transfer (Bio-Rad) for 7 min at 2.5 A. Membranes were incubated in Everyblot blocking buffer (Bio-Rad) for 1 h at RT. For Iba-1 probe, membranes were cut at 37 kDa marker and lower half was incubated with Iba1, top half was incubated with b-actin. For p-Tau, membranes were cut between 75 and 100 kDa marker. Top half was incubated with vinculin, bottom half was incubated with phospho-Tau (Thr231) from Cell Signaling (cat # 71429). All primary antibody incubations were overnight at 4°C. Membranes were washed in 1X Tris-buffered saline with 0.1% Tween-20 (TBST) prior to a secondary incubation with either goat anti-mouse HRP or goat anti-rabbit HRP at a dilution of 1:3000 for 1 h at RT. Membranes were washed a second time with 1X TBST prior to a 5 min incubation in ECL reagent (superSignal west pico plus ECL substrate kit). Blots were imaged on a Bio-Rad Chemidoc. Densitometric analysis was performed in ImageLab Software (Bio-Rad). Target signals were normalized to a control protein on each blot of either Vinculin or β-actin. Aliquots of the RIPA extracted proteins (from above) were used to quantify the cytokines by ELISA for IL-1β (Invitrogen IL-1 beta Mouse ELISA kit, cat # BMS6002), IL-6 (Invitrogen IL-6 Mouse ELISA kit, cat # BMS603-2) and TNFα (Invitrogen TNF alpha Mouse ELISA kit, cat # BMS607-3), from whole brains according to manufacturer’s protocol. All samples were run in duplicate. The average from both wells was used for quantification. Original western blots can be found in Supporting information S1 and S2.

### Nucleic Acid Extraction, PCR, and RT-qPCR

DNA and RNA was extracted for sexing embryonic mice and RT-qPCR analysis of cytokine mRNAs present in the pup brains. Nucleic acids were extracted from frozen whole brains using the Omega Bio-tek E.Z.N.A. kit according to manufacturer’s protocol (cat# R6731-01). RNA and DNA concentrations were determined using Qubit Fluorometer (Applied Biosystems) prior to subsequent analysis. For sexing of pups we added 1µL DNA to a reaction with the Q5 Master Mix (NEB, cat# M04925) and primer sets for amplification of the IL3 (For 5’-GGG ACT CCA AGC TTC AAT CA-3’, Rev 5’-TGG AGG AAG AAA AGC AA-3’) and Sry (For 5’-TGG GAC TGG TGA CAA TTG TC, Rev 5’-GAG TAC AGG TGT GCA GCT CT-3’) genes (57). PCR conditions were 95°C for 4.5 min followed by 33 cycles of 95°C for 35 s, 50°C for 1 min and 72°C for 1 min. PCR products were terminated with a final extension at 72°C for 5 min. Bands positive for both are labeled as male while samples only expressing IL3 are labeled as female.

cDNAs for RT-qPCR were made by adding 1000 ng RNA to a reaction using the High Capacity cDNA Reverse Transcription Kit (Applied Biosystems, cat# 4368814). 1µL of cDNA was added per reaction with gene specific primer pair sets for GAPDH, IL-1β, IL-6, TNFα, IL-4, IL-10, TGFβ, MyD88, NFκβ, and NLRP3 (Integrated DNA Technologies, PrimeTime qPCR Primer Assay). All samples were run in triplicate on a QuantStudio6 Pro (Applied Biosystems) using the PowerUp SYBR Green Master Mix (ThermoFisher, cat # A25741). Cycling conditions are as follows: 95°C for 15s followed by 60°C for 1 min for 40 cycles. Gene expression was measured using the QuantStudio6 Pro detection system and normalized to GAPDH. Ct values were averaged from each triplicate and used to calculate the relative expression by the 2^^-(ΔΔCt)^ method with GAPDH as the reference gene.

### Immunofluorescent Staining

Fixed brains were embedded in OCT medium and 25 µm sagittal sections were collected by a cryostat. Sections were washed with PBS, blocked in PBS containing 0.1% tritonX-100, and 10% donkey serum and probed with and primary antibodies to Cux1 (Protein Tech 11733-1-AP; diluted 1:300), SatB2 (Abcam ab92446; diluted 1:100), and Ctip2 (Abcam ab-18465; diluted 1:500). Sections were then washed with PBS and incubated with Alexa Fluor® secondary antibodies (Thermo Fisher Scientific) and DAPI followed by mounting with Prolong Diamond Antifade (Thermo Fisher Scientific). Images were acquired with an ImageXpress Micro Confocal High Content Imaging System (IXMC; Molecular Devices) and analyzed using MetaXpress software (Molecular Devices).

### Data Analysis

We used the analysis routines in GraphPad PRISM v. 10.0 to perform statistical analysis and plotting of data points. Outlier data points were identified by the ROUT method with Q = 1% and removed from analysis. Data were tested for normality with the Kolmogorov-Smirnov test. Group differences for data with normal distributions were tested with the two-tailed, unpaired Student’s-test, otherwise statistical differences were determined with the Mann Whitney-U test. Data are presented with the median and 95% Confidence Interval (CI).

## Supporting information

S1. Iba-1 and Thr231 western blots.

Reagent List

## Acknowledgments

We thank Dr. Rolf Stottmann and Dr. Tracy Bedrosian in the Institute for Genomic Medicine for their guidance on collecting and analyzing the cortical neurons of the embryonic mouse brains. We also thank Dr. Juhi Bagaitkar in the Center for Microbial Pathogenesis at Nationwide Children’s Hospital or her discussions on the immunological implications of *P. gingivalis* infection.

## Supporting Information

S1. Iba-1 and Thr231 western blots.

## References

1. Carrillo-de-Albornoz A, Figuero E, Herrera D, Bascones-Martínez A. Gingival changes during pregnancy: II. Influence of hormonal variations on the subgingival biofilm. Journal of Clinical Periodontology. 2010;37(3):230–40.

2. Fujiwara N, Tsuruda K, Iwamoto Y, Kato F, Odaki T, Yamane N, et al. Significant increase of oral bacteria in the early pregnancy period in Japanese women. Journal of Investigative and Clinical Dentistry. 2017;8(1):e12189.

3. Lin W, Jiang W, Hu X, Gao L, Ai D, Pan H, et al. Ecological Shifts of Supragingival Microbiota in Association with Pregnancy. Frontiers in Cellular and Infection Microbiology. 2018;8.

4. Paropkari AD, Leblebicioglu B, Christian LM, Kumar PS. Smoking, pregnancy and the subgingival microbiome. Sci Rep. 2016;6(1):30388.

5. Jang H, Patoine A, Wu TT, Castillo DA, Xiao J. Oral microflora and pregnancy: a systematic review and meta-analysis. Sci Rep. 2021;11(1):16870.

6. Powell AM, Khan FZA, Ravel J, Elovitz MA. Untangling Associations of Microbiomes of Pregnancy and Preterm Birth. Clinics in Perinatology. 2024;51(2):425–39.

7. Wen P, Li H, Xu X, Zhang F, Zhao D, Yu R, et al. A prospective study on maternal periodontal diseases and neonatal adverse outcomes. Acta Odontol Scand. 2024;83:348–55.

8. Kassebaum NJ, Bernabé E, Dahiya M, Bhandari B, Murray CJL, Marcenes W. Global Burden of Severe Periodontitis in 1990-2010:A Systematic Review and Meta-regression. Journal of Dental Research. 2014;93(11):1045–53.

9. Lieff S, Boggess KA, Murtha AP, Jared H, Madianos PN, Moss K, et al. The Oral Conditions and Pregnancy Study: Periodontal Status of a Cohort of Pregnant Women. Journal of Periodontology. 2004;75(1):116–26.

10. Kornman KS, Loesche WJ. The subgingival microbial flora during pregnancy. J Periodontal Res. 1980;15(2):111–22.

11. Reyes L, Phillips P, Wolfe B, Golos TG, Walkenhorst M, Progulske-Fox A, et al. Porphyromonas gingivalis and adverse pregnancy outcome. J Oral Microbiol. 2018;10(1):1374153.

12. Silva de Araujo Figueiredo C, Gonçalves Carvalho Rosalem C, Costa Cantanhede AL, Abreu Fonseca Thomaz ÉB, Fontoura Nogueira da Cruz MC. Systemic alterations and their oral manifestations in pregnant women. Journal of Obstetrics and Gynaecology Research. 2017;43(1):16-22.

13. Chaparro A, Blanlot C, Ramirez V, Sanz A, Quintero A, Inostroza C, et al. Porphyromonas gingivalis, Treponema denticola and toll-like receptor 2 are associated with hypertensive disorders in placental tissue: a case-control study. J Periodontal Res. 2013;48(6):802–9.

14. Ercan E, Eratalay K, Deren O, Gur D, Ozyuncu O, Altun B, et al. Evaluation of periodontal pathogens in amniotic fluid and the role of periodontal disease in pre-term birth and low birth weight. Acta Odontol Scand. 2013;71(3-4):553–9.

15. Gonzales-Marin C, Spratt DA, Millar MR, Simmonds M, Kempley ST, Allaker RP. Levels of periodontal pathogens in neonatal gastric aspirates and possible maternal sites of origin. Mol Oral Microbiol. 2011;26(5):277–90.

16. Katz J, Chegini N, Shiverick KT, Lamont RJ. Localization of P. gingivalis in preterm delivery placenta. J Dent Res. 2009;88(6):575–8.

17. Leon R, Silva N, Ovalle A, Chaparro A, Ahumada A, Gajardo M, et al. Detection of Porphyromonas gingivalis in the amniotic fluid in pregnant women with a diagnosis of threatened premature labor. J Periodontol. 2007;78(7):1249–55.

18. Swati P, Ambika Devi K, Thomas B, Vahab SA, Kapaettu S, Kushtagi P. Simultaneous detection of periodontal pathogens in subgingival plaque and placenta of women with hypertension in pregnancy. Arch Gynecol Obstet. 2012;285(3):613–9.

19. Ishida E, Furusho H, Renn T-Y, Shiba F, Chang H-M, Oue H, et al. Mouse maternal odontogenic infection with Porphyromonas gingivalis induces cognitive decline in offspring. Frontiers in Pediatrics. 2023;11.

20. Lara B, Loureiro I, Gliosca L, Castagnola L, Merech F, Gallino L, et al. Porphyromonas gingivalis outer membrane vesicles shape trophoblast cell metabolism impairing functions associated to adverse pregnancy outcome. Journal of Cellular Physiology. 2023;238(11):2679–91.

21. Ao M, Miyauchi M, Furusho H, Inubushi T, Kitagawa M, Nagasaki A, et al. Dental Infection of Porphyromonas gingivalis Induces Preterm Birth in Mice. PLOS ONE. 2015;10(8):e0137249.

22. Konishi H, Urabe S, Miyoshi H, Teraoka Y, Maki T, Furusho H, et al. Fetal Membrane Inflammation Induces Preterm Birth Via Toll-Like Receptor 2 in Mice With Chronic Gingivitis. Reprod Sci. 2019;26(7):869–78.

23. Konishi H, Urabe S, Teraoka Y, Morishita Y, Koh I, Sugimoto J, et al. Porphyromonas gingivalis, a cause of preterm birth in mice, induces an inflammatory response in human amnion mesenchymal cells but not epithelial cells. Placenta. 2020;99:21–6.

24. Lin D, Smith MA, Champagne C, Elter J, Beck J, Offenbacher S. Porphyromonas gingivalis infection during pregnancy increases maternal tumor necrosis factor alpha, suppresses maternal interleukin-10, and enhances fetal growth restriction and resorption in mice. Infect Immun. 2003;71(9):5156–62.

25. Yoshida S, Hatasa M, Ohsugi Y, Tsuchiya Y, Liu A, Niimi H, et al. Porphyromonas gingivalis Administration Induces Gestational Obesity, Alters Gene Expression in the Liver and Brown Adipose Tissue in Pregnant Mice, and Causes Underweight in Fetuses. Frontiers in Cellular and Infection Microbiology. 2022;11.

26. Choi J-W, Kim S-C, Hong S-H, Lee H-J. Secretable Small RNAs via Outer Membrane Vesicles in Periodontal Pathogens. Journal of Dental Research. 2017;96(4):458–66.

27. Gui MJ, Dashper SG, Slakeski N, Chen Y-Y, Reynolds EC. Spheres of influence: Porphyromonas gingivalis outer membrane vesicles. Molecular Oral Microbiology. 2016;31(5):365–78.

28. Ho M-H, Chen C-H, Goodwin JS, Wang B-Y, Xie H. Functional Advantages of Porphyromonas gingivalis Vesicles. PLOS ONE. 2015;10(4):e0123448.

29. Okamura H, Hirota K, Yoshida K, Weng Y, He Y, Shiotsu N, et al. Outer membrane vesicles of Porphyromonas gingivalis: Novel communication tool and strategy. Japanese Dental Science Review. 2021;57:138–46.

30. Veith PD, Chen Y-Y, Gorasia DG, Chen D, Glew MD, O’Brien-Simpson NM, et al. Porphyromonas gingivalis Outer Membrane Vesicles Exclusively Contain Outer Membrane and Periplasmic Proteins and Carry a Cargo Enriched with Virulence Factors. Journal of Proteome Research. 2014;13(5):2420–32.

31. Zhang J, Yu C, Zhang X, Chen H, Dong J, Lu W, et al. Porphyromonas gingivalis lipopolysaccharide induces cognitive dysfunction, mediated by neuronal inflammation via activation of the TLR4 signaling pathway in C57BL/6 mice. J Neuroinflammation. 2018;15(1):37.

32. Nara PL, Sindelar D, Penn MS, Potempa J, Griffin WST. Porphyromonas gingivalis Outer Membrane Vesicles as the Major Driver of and Explanation for Neuropathogenesis, the Cholinergic Hypothesis, Iron Dyshomeostasis, and Salivary Lactoferrin in Alzheimer’s Disease. J Alzheimers Dis. 2021;82(4):1417–50.

33. Gong T, Chen Q, Mao H, Zhang Y, Ren H, Xu M, et al. Outer membrane vesicles of Porphyromonas gingivalis trigger NLRP3 inflammasome and induce neuroinflammation, tau phosphorylation, and memory dysfunction in mice. Front Cell Infect Microbiol. 2022;12:925435.

34. Lan C, Chen S, Jiang S, Lei H, Cai Z, Huang X. Different expression patterns of inflammatory cytokines induced by lipopolysaccharides from Escherichia coli or Porphyromonas gingivalis in human dental pulp stem cells. BMC Oral Health. 2022;22(1):121.

35. Qiu C, Yuan Z, He Z, Chen H, Liao Y, Li S, et al. Lipopolysaccharide Preparation Derived From Porphyromonas gingivalis Induces a Weaker Immuno-Inflammatory Response in BV-2 Microglial Cells Than Escherichia coli by Differentially Activating TLR2/4-Mediated NF-κB/STAT3 Signaling Pathways. Frontiers in Cellular and Infection Microbiology. 2021;11.

36. Bodet C, Chandad F, Grenier D. Modulation of cytokine production by Porphyromonas gingivalis in a macrophage and epithelial cell co-culture model. Microbes and Infection. 2005;7(3):448–56.

37. Castillo Y, Castellanos JE, Lafaurie GI, Castillo DM. Porphyromonas gingivalis outer membrane vesicles modulate cytokine and chemokine production by gingipain-dependent mechanisms in human macrophages. Archives of Oral Biology. 2022;140:105453.

38. Charoensaensuk V, Chen YC, Lin YH, Ou KL, Yang LY, Lu DY. Porphyromonas gingivalis Induces Proinflammatory Cytokine Expression Leading to Apoptotic Death through the Oxidative Stress/NF-κB Pathway in Brain Endothelial Cells. Cells. 2021;10(11).

39. Palm E, Khalaf H, Bengtsson T. Porphyromonas gingivalis downregulates the immune response of fibroblasts. BMC Microbiology. 2013;13(1):155.

40. Yang K, Xu S, Zhao H, Liu L, Lv X, Hu F, et al. Hypoxia and Porphyromonas gingivalis-lipopolysaccharide synergistically induce NLRP3 inflammasome activation in human gingival fibroblasts. International Immunopharmacology. 2021;94:107456.

41. Offenbacher S, Katz V, Fertik G, Collins J, Boyd D, Maynor G, et al. Periodontal infection as a possible risk factor for preterm low birth weight. J Periodontol. 1996;67(10 Suppl):1103-13.

42. Zhang B, Wei YZ, Wang GQ, Li DD, Shi JS, Zhang F. Targeting MAPK Pathways by Naringenin Modulates Microglia M1/M2 Polarization in Lipopolysaccharide-Stimulated Cultures. Front Cell Neurosci. 2018;12:531.

43. Frank-Cannon TC, Alto LT, McAlpine FE, Tansey MG. Does neuroinflammation fan the flame in neurodegenerative diseases? Mol Neurodegener. 2009;4:47.

44. Gao Y, Tan L, Yu JT, Tan L. Tau in Alzheimer’s Disease: Mechanisms and Therapeutic Strategies. Curr Alzheimer Res. 2018;15(3):283–300.

45. Morris SL, Brady ST. Tau phosphorylation and PAD exposure in regulation of axonal growth. Frontiers in Cell and Developmental Biology. 2023;10.

46. Seubert P, Mawal-Dewan M, Barbour R, Jakes R, Goedert M, Johnson GV, et al. Detection of phosphorylated Ser262 in fetal tau, adult tau, and paired helical filament tau. J Biol Chem. 1995;270(32):18917–22.

47. Brion J-P, Smith C, Couck A-M, Gallo J-M, Anderton BH. Developmental Changes in τ Phosphorylation: Fetal τ Is Transiently Phosphorylated in a Manner Similar to Paired Helical Filament-τ Characteristic of Alzheimer’s Disease. Journal of Neurochemistry. 1993;61(6):2071–80.

48. Tang Z, Liang D, Cheng M, Su X, Liu R, Zhang Y, et al. Effects of Porphyromonas gingivalis and Its Underlying Mechanisms on Alzheimer-Like Tau Hyperphosphorylation in Sprague-Dawley Rats. Journal of Molecular Neuroscience. 2021;71(1):89–100.

49. Yu Y, Run X, Liang Z, Li Y, Liu F, Liu Y, et al. Developmental regulation of tau phosphorylation, tau kinases, and tau phosphatases. J Neurochem. 2009;108(6):1480–94.

50. Arendt T, Stieler JT, Holzer M. Tau and tauopathies. Brain Res Bull. 2016;126(Pt 3):238–92.

51. Wang C, Fan L, Khawaja RR, Liu B, Zhan L, Kodama L, et al. Microglial NF-κB drives tau spreading and toxicity in a mouse model of tauopathy. Nature Communications 2015 6. 2022;13(1):1969.

52. Perea JR, Ávila J, Bolós M. Dephosphorylated rather than hyperphosphorylated Tau triggers a pro-inflammatory profile in microglia through the p38 MAPK pathway. Experimental Neurology. 2018;310:14–21.

53. Hefti MM, Kim S, Bell AJ, Betters RK, Fiock KL, Iida MA, et al. Tau Phosphorylation and Aggregation in the Developing Human Brain. J Neuropathol Exp Neurol. 2019;78(10):930–8.

54. Sapir T, Frotscher M, Levy T, Mandelkow EM, Reiner O. Tau’s role in the developing brain: implications for intellectual disability. Hum Mol Genet. 2012;21(8):1681–92.

55. Mishra A, Bandopadhyay R, Singh PK, Mishra PS, Sharma N, Khurana N. Neuroinflammation in neurological disorders: pharmacotherapeutic targets from bench to bedside. Metab Brain Dis. 2021;36(7):1591–626.

56. Mashburn LM, Whiteley M. Membrane vesicles traffic signals and facilitate group activities in a prokaryote. Nature. 2005;437(7057):422-5.

57. Lambert J-F, Benoit BO, Colvin GA, Carlson J, Delville Y, Quesenberry PJ. Quick sex determination of mouse fetuses. Journal of Neuroscience Methods. 2000;95(2):127–32.

