## Supplementary material for "*Porphyromonas gingivalis* outer membrane vesicles alter neuronal architecture and Tau phosphorylation in the embryonic mouse brain": S1. Iba-1 and Thr231 western blots.

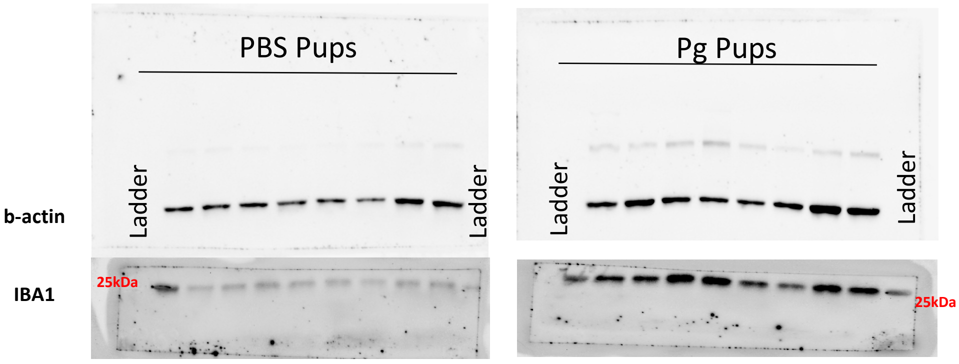


**Supporting Fig 1:**
Full western blot images from pup brain lysates probed with IBA1 and b-actin antibodies.

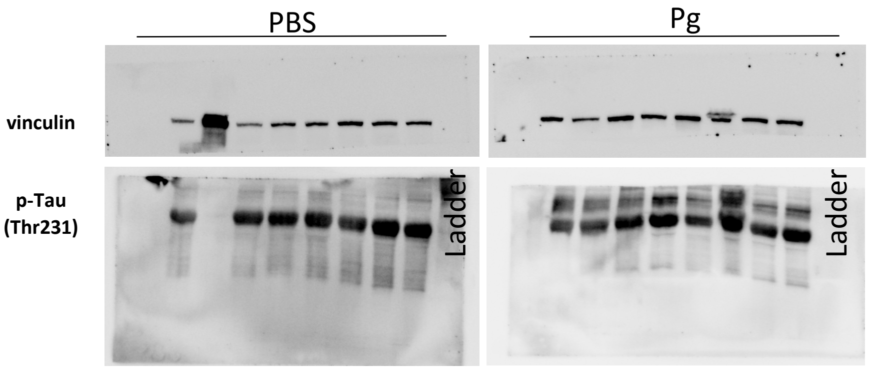


**Supporting Fig 2:** Full western blot images from pup brain lysates probed with pTau(Thr231) and vinculin antibodies
